# The human brain networks mediating the vestibular sensation of self-motion

**DOI:** 10.1101/2021.12.03.471139

**Authors:** Zaeem Hadi, Yuscah Pondeca, Elena Calzolari, Mohammad Mahmud, Mariya Chepisheva, Rebecca M Smith, Heiko Rust, David J Sharp, Barry M Seemungal

## Abstract

Vestibular Agnosia - where peripheral vestibular activation triggers the usual reflex nystagmus response but with attenuated or no self-motion perception - is found in brain disease with disrupted cortical network functioning, e.g. traumatic brain injury (TBI) or neurodegeneration (Parkinson’s Disease). Patients with acute focal hemispheric lesions (e.g. stroke) do not manifest vestibular agnosia. Thus brain network mapping techniques, e.g. resting state functional MRI (rsfMRI), are needed to interrogate functional brain networks mediating vestibular agnosia. Whole-brain rsfMRI was acquired from 39 prospectively recruited acute TBI patients with preserved peripheral vestibular function, along with self-motion perceptual thresholds during passive yaw rotations in the dark. Following quality-control checks, 25 patient scans were analyzed. TBI patients were classified as having vestibular agnosia (n = 11) or not (n = 14) via laboratory testing of self-motion perception. Using independent component analysis, we found altered functional connectivity in the right superior longitudinal fasciculus and left rostral prefrontal cortex in vestibular agnosia. Moreover, regions of interest analyses showed both inter-hemispheric and intra-hemispheric network disruption in vestibular agnosia. In conclusion, our results show that vestibular agnosia is mediated by bilateral anterior and posterior network dysfunction and reveal the distributed brain mechanisms mediating vestibular self-motion perception.

## 1 Introduction

We recently characterized a clinical syndrome called vestibular agnosia (VA) in patients with acute traumatic brain injury (TBI), where, despite preserved peripheral and reflex vestibular functioning, patients had an attenuated vestibular-mediated sensation of self-motion (‘vestibular-motion perception’) (Calzolari et al., 2021). Vestibular agnosia has also been recorded in elderly patients, typically with small vessel disease and imbalance (Chiarovano et al., 2016; Imbaud Genieys, 2007; Seemungal, 2006), and in patients with advanced Parkinson’s Disease with falls (Yousif et al., 2016). We recently showed that in acute TBI patients with imbalance, objective measures of VA correlate with impaired white matter structural integrity in the right inferior longitudinal fasciculus (ILF) (Calzolari et al., 2021). This structural analysis thus identified the overlap in imbalance and VA, however the neural correlates distinct to VA were not identified.

There is evidence to support the notion that self-motion perception is not localizable. We previously showed that the duration of motion-perception sensation in the dark (elicited by a rapid stop from constant angular rotation) in healthy individuals correlated with a widespread, bilateral white matter network (Nigmatullina et al., 2015). Subsequent tractography studies also showed that vestibular networks are bihemispheric linked via corpus callosum (Kirsch et al., 2016; Wirth et al., 2018). In support of a distributed coding for self-motion perception is the lack of effect of acute focal lesions (stroke - (Kaski et al., 2016)) or via non-invasive stimulation (Seemungal et al., 2008) on vestibular-motion perception.

Generally, the posterior brain networks are considered to encode sensory inputs whereas anterior networks encode higher order functions and integration of sensory inputs (Ester et al., 2016). This large-scale coordination of brain networks has been shown in spatial and visual sensory modalities in primates and humans, where sensory processing and integration is associated with unique activation within parietal regions (intra-network) and a distinct activation of fronto-parietal (inter-) networks (Dotson et al., 2014; Fiebelkorn et al., 2018; Galati et al., 2001). Similar unique activations associated with anterior integration network, posterior sensory processing networks, and large scale anterior-posterior inter-network connectivity has been linked to cross-modal sensory perception disorders (e.g. synesthesia, Dovern et al., 2012). Accordingly, we postulated that there are (at least) two main networks mediating VA; a posterior brain network, receiving the main bottom-up vestibular signals mediating sensory processing, and an anterior network linked to sensory integration and perceptual ignition (Del Cul et al., 2007). Thus, dysfunction of the posterior network could result in loss of perception via impaired afferent signal processing; whereas a sensory integration dysfunction in anterior networks could also mediate VA.

Currently no study has shown the brain regions specifically associated with impaired self-motion perception with intact peripheral functioning (i.e. VA). Our general objective was to identify brain regions associated with VA. Moreover, to test our postulate of an anterior-posterior cortical network mediating VA, we hypothesize that VA in acute TBI is mediated by multi-network dysfunction, i.e. (i) by impaired functional connectivity within anterior networks, (ii) impaired functional connectivity within posterior networks, and (iii) inter-network impaired functional connectivity. We further hypothesize that posterior networks containing putative vestibular regions (parieto-insular vestibular cortex, mid temporal regions) will have disrupted functional projections. Finally, since we previously showed that VA was linked to imbalance via damage to the right inferior longitudinal fasciculus (ILF) (Calzolari et al., 2021) and ILF connects higher order visual cortices to the temporal regions (Latini et al., 2017; Panesar et al., 2018), we probed the cortical regions mapped by right ILF using the lingual gyrus as a seed, a region often linked to motion- and illusion-perception (Cha & Chakrapani, 2015; Della-Justina et al., 2014; Kahane et al., 2003; Roberts et al., 2017). To assess our predictions, we evaluated functional connectivity via resting state fMRI in both grey and white matter brain networks.

## 2 Material and Methods

### 2.1 Participants and Recruitment

The data reported in this paper was collected as part of an MRC-funded prospective study (Calzolari et al., 2021). 146 acute TBI patients were clinically assessed from whom 39 were recruited from the St Mary’s Hospital Major Trauma Centre (London, UK) and King’s College Hospital (London, UK). Inclusion Criteria were: (i) blunt head injury resulting in admission to the major trauma ward; (ii) age 18-65; (iii) preserved vestibular function. Exclusion criteria were: (i) additional active pre-morbid medical, neurological, or psychiatric condition (unless inactive or controlled); (ii) musculoskeletal condition impairing ability to balance; (iii) substance abuse history; (iv) pregnancy; and (v) inability to obtain consent or assent. Thirty-seven matched healthy controls were also recruited following written informed consent. The study was conducted in accordance with the principles of the Declaration of Helsinki and was approved by the local Research Ethics Committee.

### 2.2 Procedure

All participants completed assessment of peripheral and reflex vestibular function, vestibular perceptual testing, reaction times, posturography, neuroimaging, and questionnaires for perceived balance, and cognitive examination. The details for all tests has been reported previously (Calzolari et al., 2021). Procedures involved in assessment of peripheral vestibular dysfunction, vestibular perceptual testing, posturography, and resting-state scans are listed here.

### 2.3 Assessment of Peripheral Vestibular Function

Patients were assessed for their peripheral vestibular function to exclude peripheral dysfunction as a cause of impaired vestibular perception. Video head impulse testing and rotational chair testing with eye movement assessment of VOR gain for the stopping response from 90°/s constant rotation was used to assess peripheral dysfunction. All TBI patients included in the study had intact peripheral vestibular function (Calzolari et al., 2021).

### 2.4 Vestibular Perceptual Threshold

Vestibular perceptual thresholds were determined as a measure of self-motion perception and to classify subjects having abnormal perceptual thresholds with VA (Calzolari et al., 2021). Figure 1 shows the apparatus and method used to determine participants’ vestibular-perceptual thresholds during passive yaw-plane rotations in the dark (Seemungal et al., 2004). Participants sat on a rotating chair in dark and were instructed to press right or left button as soon as they perceive the rotation in respective direction (Figure 1A and Figure 1B). Lights were turned on after each trial in order to allow the decay of post-rotatory vestibular effects. White noise was provided via 2 speakers (for each ear) attached with the chair to mask any auditory cues from the environment.

**Figure 1.**
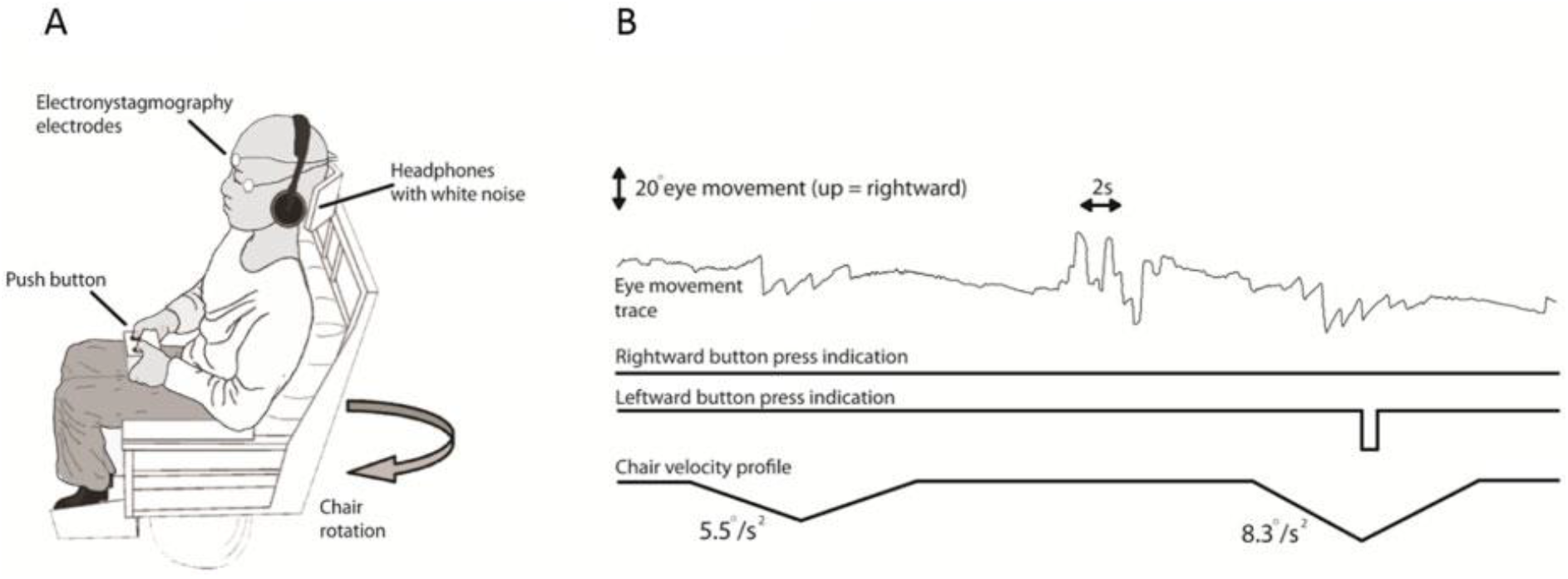
Vestibular thresholds. Apparatus and methods. **(A)** Rotating Chair. **(B)** Raw traces for two subsequent rotations for a patient.

### 2.5 Posturography

Postural sway was assessed using a force platform for 60 seconds duration under four conditions: hard surface with eyes open (HO), hard surface with eyes closed (HC), soft surface with eyes open (SO), and soft surface with eyes closed (SC). Subjects were instructed to stand with their arms hanging loosely and to maintain their balance. SC condition is primarily vestibular dependent due to reduced proprioceptive (soft surface) and visual feedback (eyes closed); SC has also been shown to be the condition that best predicts difference in balance between TBI patients and healthy controls (Calzolari et al., 2021). Root mean square (RMS) sway calculated using custom MATLAB scripts, during SC condition was thus chosen as the covariate for removing the confounding effect of balance from motion perception in neuroimaging analysis.

### 2.6 Vestibular Agnosia Classification

Vestibular perceptual thresholds from 37 healthy participants (Mean Perceptual Thresholds: 0.76 deg/s^2^, SD = 0.42) were used to establish a normative range. We have previously used a more conservative approach by classifying patients with having VA if their mean perceptual threshold (average of thresholds in right and left direction rotations) were above the mean + 3 standard deviation of the perceptual thresholds of healthy controls (Calzolari et al., 2021). This criterion enhances specificity but at the cost of reduced sensitivity. We thus used a slightly lenient and arguably objective approach for classification using all 76 subjects (healthy: 37; TBI: 39) through k-means clustering. We identified 3 clusters (Figure 2): i) cluster 1 was composed of subjects with normal perceptual thresholds (labeled as healthy controls and normal TBI in Figure 2); ii) cluster 2 was composed of subjects with moderately high perceptual thresholds (labeled as healthy outliers and moderate VA TBI in Figure 2), and iii) cluster 3 was composed of subjects with high perceptual thresholds (labeled as high VA TBI in Figure 2). 19 TBI patients from cluster 2 and 3 with moderately high and high perceptual thresholds respectively (Mean Perceptual Thresholds: 4.20 deg/s^2^, SD = 3.92), were classified as having vestibular agnosia (VA+) whereas 20 TBI patients (Mean Perceptual Thresholds: 0.85 deg/s^2^, SD = 0.28) clustered with the healthy subjects in cluster 1 were considered patients without vestibular agnosia (VA-). Healthy control data was not used in the imaging analysis (and were used only in supporting the classification of vestibular agnosia). Imaging analysis compared TBI patients without vestibular agnosia (VA-) who were considered the control group, with TBI patients with vestibular agnosia (VA+).

**Figure 2.**
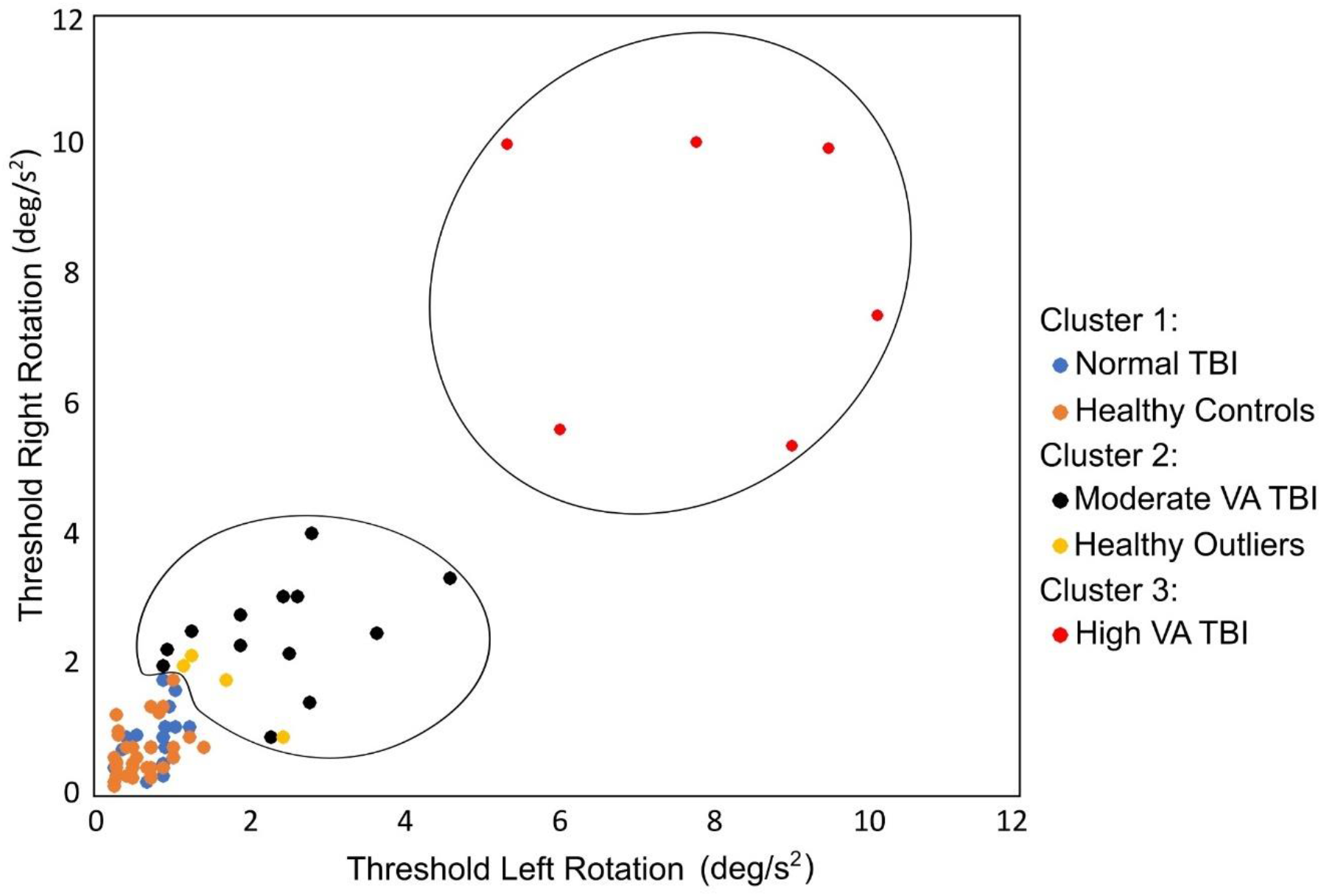
K-means clustering with 3 clusters. K-mean clustering was performed on subjects’ perceptual thresholds on right and left sides and three clusters were determined.

## 3 Neuroimaging

Historically, fMRI studies have assessed activity in grey matter (Ogawa et al., 1992) whereas fMRI activity within the white-matter was considered noise. However, recent evidence suggests that the BOLD signal in white matter has similar hemodynamic response characteristics as in grey matter (Fraser et al., 2012) and the functional activity in underlying white-matter tracts maps along the direction of white-matter tracts (Ding et al., 2013, 2016). Several studies have reported the presence of a white-matter functional architecture corresponding to resting state functional activity (Huang et al., 2020; Li et al., 2020; Peer et al., 2017) as well as task related activity (Huang et al., 2018, 2020; Marussich et al., 2017).

We thus performed neuroimaging analysis in grey- and white-matter regions separately using resting-state fMRI scans. After excluding 14 of 39 TBI patients due to different field of view scanning parameters (n =12 of 39), excessive motion artifacts (n =1 of 39), and failed segmentation (n = 1 of 39), we were left with 25 TBI patients (11 VA+ and 14 VA-) whose data were included in the neuroimaging analysis.

### 3.1 Image Acquisition

Structural and functional MRI images were acquired using a 3T Siemens Verio (Siemens) scanner, using a 32-channel head coil. The scanning protocol included: (i) 3D T1-weighted images acquired using MPRAGE sequence (image matrix: 256 x 256; voxel size: 1 x 1; Slices: 160; field of view: 256 x 256 mm; slice thickness: 3 mm; TR = 2300 ms; TE: 2.98 ms); and (ii) T2*-weighted images sensitive to blood oxygenation level dependent (BOLD) signal for resting state fMRI (image matrix: 64 x 64; voxel size: 3 x 3 x 3 mm^3^; Slices: 35; field of view: 192 x 192 mm; flip angle: 80°; slice thickness: 3 mm; TR = 2000 ms; TE: 30 ms; volumes = 300; scan time = 10 minutes). Subjects were instructed to keep their eyes closed, stay awake, and to try not to think of anything.

### 3.2 Preprocessing

Data were preprocessed using the CONN Toolbox (Whitfield-Gabrieli & Nieto-Castanon, 2012) based on Statistical Parametric Mapping (SPM12; http://www.fil.ion.ucl.ac.uk/spm/). Preprocessing steps were as follows. (1) Realignment to mean functional image, unwarping, and susceptibility distortion correction. (2) Slice timing correction. (3) Functional outlier detection using ART (artifact detection toolbox), with scans exceeding framewise displacement of 0.5 mm were labeled as outliers (Power et al., 2012). (4) Structural segmentation and normalization. (5) Functional normalization using deformation fields estimated from structural normalization. (6) Semi-automated lesion masking using T1 images with ITK-SNAP software (Yushkevich et al., 2006) and subtraction of lesions from individual participants’ segmented maps. (7) Group analysis space masks: two analysis space masks were created: 1) grey-matter (GM) specific; 2) white-matter (WM) specific. Segmented masks of individual participants were intersected with functional masks to remove regions with missing functional data. These individual masks were binarized (GM: p > 0.2; WM: p > 0.8) and then averaged across subjects. Voxels identified in >70% subjects were used for GM specific mask creation whereas voxels in >90% were selected for WM specific mask creation. (8) Lesion masks of all subjects were then combined and subtracted from the two group analysis space masks. (9) Smoothing (6 mm FWHM) was performed within the grey- and white-matter using GM and WM masks separately. (10) Subsequently, denoising was performed using aCompCor (Behzadi et al., 2007) approach as implemented in CONN, in which six motion regressors (3 translational and 3 angular motion), their temporal derivatives, cerebrospinal fluid (10 principal components), and outlier scans identified by ART toolbox were regressed out. One subject with >50% (199 of 300) outlier scans was removed from the analysis. (11) Data was then band-pass filtered with frequency range of 0.008-0.1 Hz. White matter activity and the global signal were not regressed out since white-matter activity was the signal of interest whereas the global signal regression is known to introduce negative correlations (Murphy et al., 2009).

### 3.3 Second-Level Analysis

Group level analysis was performed to determine the differences between VA+ (n = 11) TBI patients and VA- (n = 14) TBI patients. RMS sway and lesion volume were added as covariates in group-analysis to remove the effects explained by balance and extent of injury. In addition, volume regressors were also added to control for tissue atrophy as a result of injury. Grey matter volume was added as a covariate in GM specific whereas white matter volume was added in WM specific analysis.

#### 3.3.1 Independent Component Analysis

Group ICA was performed to assess the intra-network resting state differences in VA+ and VA- groups using Fast ICA algorithm in CONN toolbox (Whitfield-Gabrieli & Nieto-Castanon, 2012). The optimal number of the independent components (ICs) were estimated in the GIFT toolbox (https://trendscenter.org/software/) using a modified minimum description length algorithm (MDL). The optimal number of ICs for GM specific analysis were found to be 38 and 10 for WM specific analysis.

In GM specific ICA, independent components containing putative vestibular regions (i.e. parietal operculum (OP2), temporo-parietal, mid temporal, thalamus, and insular regions) were considered components of interest and were evaluated for group comparisons (VA+ vs VA-). All 10 independent components were evaluated for group comparisons in WM specific ICA.

#### 3.3.2 Region of Interest Analysis

ROI Analysis was used to assess the inter-network resting state differences between VA+ and VA- groups. To select the network ROIs, we used the dice coefficient, which is a measure of spatial overlap between two images. The dice coefficient was measured for overlap between standard atlases and the independent components from our GM- and WM-specific ICA. For GM specific ROI analysis, we used an atlas of resting state networks available with CONN toolbox, determined using Human Connectome Project dataset of 497 subjects. Resting state network ROIs with dice-coefficient ≥ 0.1 (25 ROIs) were included in GM specific ROI analysis. Grey-matter resting state networks have been shown to have the dice-coefficients greater than 0.1 (Branco et al., 2016, 2020; Tie et al., 2014).

Since there are no established white-matter resting-state networks, white matter regions from the ICBM-DTI-81 atlas (Mori et al., 2008; Oishi et al., 2008) were used as ROIs. The regions with the dice-coefficient ≥ 0.01 (29 ROIs) were included in WM specific ROI analysis. As only 10 independent components were estimated in WM specific ICA, the components were not sufficient to resolve all white-matter regions available in the ICBM-DTI-81 atlas with a considerable value of the dice-coefficient. Thus, we used a lenient cut-off for the dice-coefficient in WM specific ROI analysis to include more white-matter regions in the analysis.

#### 3.3.3 Seed-Based Analysis

For testing the hypothesis that posterior networks containing putative vestibular regions will have disrupted functional projections, we selected 3 resting-state networks determined via GM specific ICA as seeds (supplementary Figure S1). All three networks contained regions from PIVC (parietal operculum), mid temporal, and insular regions. Furthermore, we probed the brain regions mapped by ILF using bilateral lingual gyri as seeds using default atlas of CONN, which is a combination of the Harvard-Oxford atlas and the AAL atlas.

## 4 Statistical Analysis

The assumptions of gaussian random field theory (RFT) are required for whole brain analysis (Nichols, 2012) and thus parametric RFT statistics are not applicable for GM or WM specific analysis. Permutation statistics (non-parametric), however, does not require gaussian RFT assumptions (Zhang et al., 2009). Moreover, the choice of cluster height threshold (either p<0.01 or p<0.001) has no impact on false positive rate when using permutation statistics for whole brain analysis as demonstrated in a previous study (Eklund et al., 2019).

Thus, all group level differences in ICA and seed-based analysis were evaluated using permutation statistics (Bullmore et al., 1999) and cluster mass FDR correction as implemented in the CONN toolbox. Cluster height threshold for all grey-matter specific analysis was selected as p<0.001 and for all white-matter specific analysis as p<0.01, as choice of height thresholds reveal different architecture in grey- and white-matter.

For WM specific ROI analysis, a connection level threshold of p<0.01 and ROI level FDR mass correction was used. For GM specific ROI analysis, we used a parametric multivariate pattern analytic (MVPA) measure called “Functional Network Connectivity” (Jafri et al., 2008) implemented in the CONN toolbox.

### 4.1 Sample Size

Effect sizes in TBI cohort are often large such that the differences between patients are easily visible to the naked eye during the clinical assessments. In our cohort, vestibular agnosia was not only apparent during clinical assessment (Calzolari et al., 2021), the difference is also very large when comparing objectively assessed perceptual thresholds of VA+ (4.20 deg/s^2^) vs VA- (0.85 deg/s^2^) with an effect size of Cohen’s d = 1.2. Considering that this is the first study looking at the functional brain correlates of VA, it was not possible to appropriately power the study. However, from previous studies, it is evident that even small sample sizes in TBI cohorts, such as 11 TBI with post-traumatic amnesia (PTA) vs 8 TBI without PTA (De Simoni et al., 2016), 12 TBI with post-concussion syndrome (PCS) vs 16 TBI without PCS (Sours et al., 2015), and 8 concussed TBI vs 11 healthy (Zhu et al., 2015) are adequate to identify functional brain differences with decent effect sizes (Hedge’s g up to 1.8). Similarly, functional brain differences in a group of 7 vs 7 TBI patients with impaired or normal balance, which is also mediated by vestibular networks, have also recently been identified (Woytowicz et al., 2018). We thus think that our analysis is adequately powered and will allow future sample size calculations for studies evaluating vestibular agnosia.

## 5 Results

### 5.1 Independent Component Analysis

#### 5.1.1 GM Specific Independent Component Analysis

Group differences were identified in IC-8, a resting-state network largely composed of bilateral superior and mid temporal regions of posterior division and of temporooccipital part (Figure 3A). VA+ group compared to VA- group had decreased functional connectivity at right intra-calcarine cortex (MNI coordinates: 06, −72, 12) in IC-8, shown in Figure 3B (t(20) = −4.42, pFDR < 0.05; effect size = −3.79).

**Figure 3.**
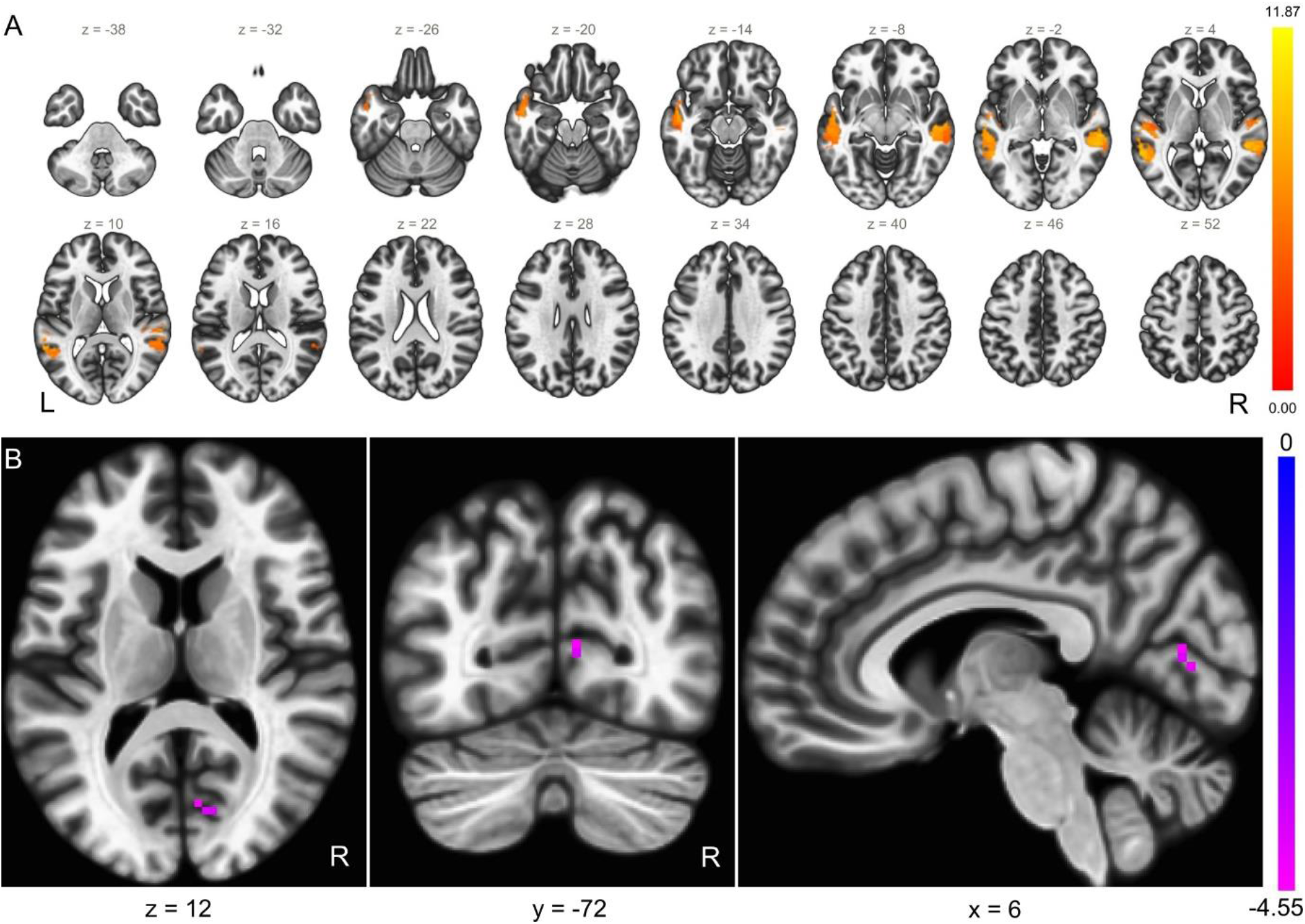
GM Specific ICA. **(A)** Independent component 8. **(B)** Group comparison between VA+ and VA- patients showing decreased functional connectivity in right intra-calcarine cortex inside the independent component-8.

Additionally, group differences were also identified in IC-18, a resting state-network composed of regions from bilateral rostral prefrontal cortex, bilateral hypothalamus and thalamus (Figure 4A). VA+ group compared to VA- group had increased functional connectivity at left frontal pole (MNI coordinates: −42, 48, 00) in IC-18, shown in Figure 4B (t(20) = 5.43, pFDR < 0.05; effect size = 4.582).

**Figure 4.**
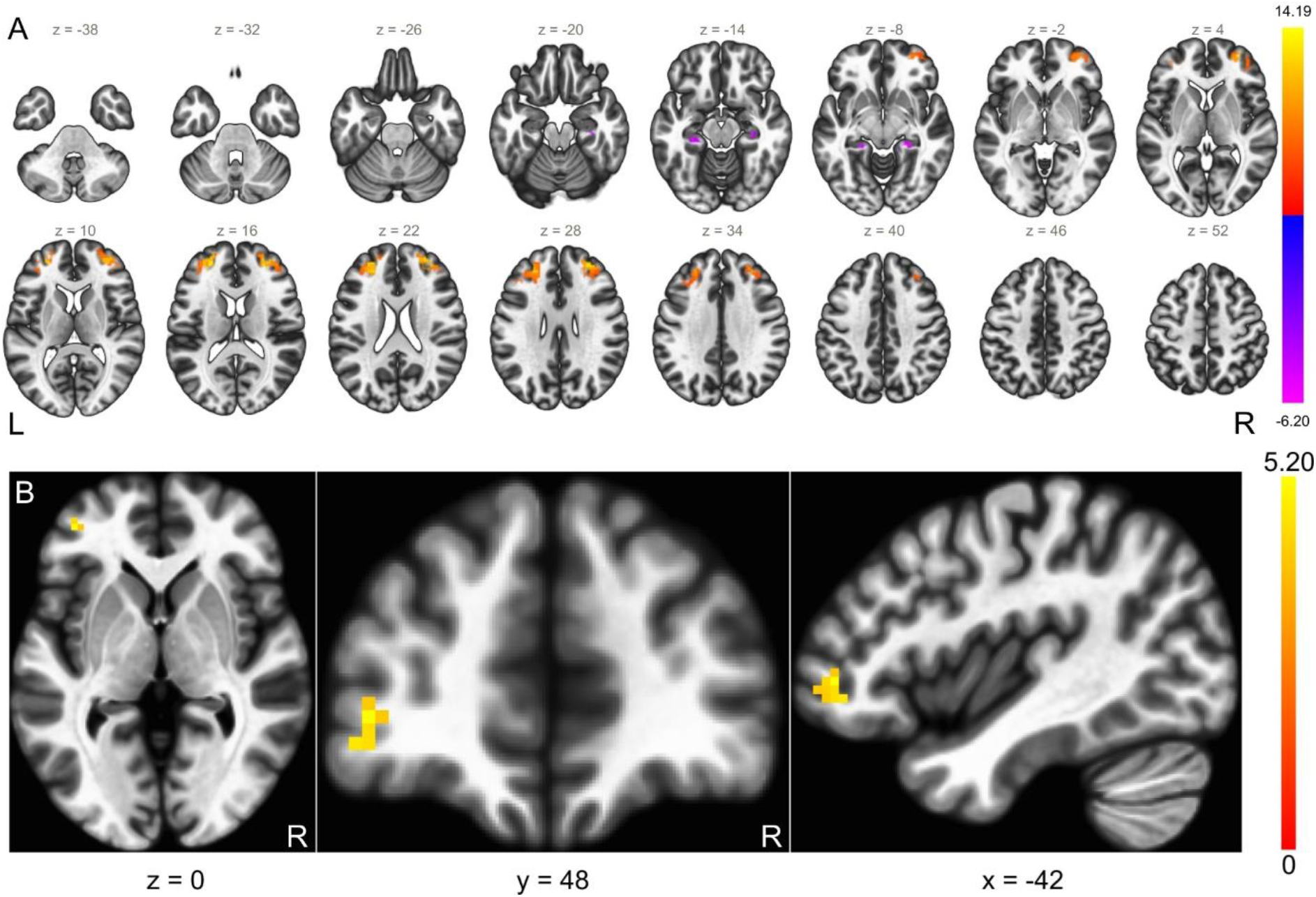
GM Specific ICA. **(A)** Independent component 18. **(B)** Group comparison between VA+ and VA- patients showing increased functional connectivity at left frontal pole inside the independent component-18.

#### 5.1.2 WM Specific Independent Component Analysis

In IC-8 (Figure 5A), we found that VA+ compared to VA- group had increased functional connectivity (t(20) = 6.34, pFDR < 0.05; effect size = 1.629) in a cluster (MNI: 39, −51, 06) composed of right superior longitudinal fasciculus (SLF), right posterior thalamic radiation (PTR), right retrolenticular part of internal capsule (RLIC), and right sagittal stratum (SS), shown in (Figure 5B).

**Figure 5.**
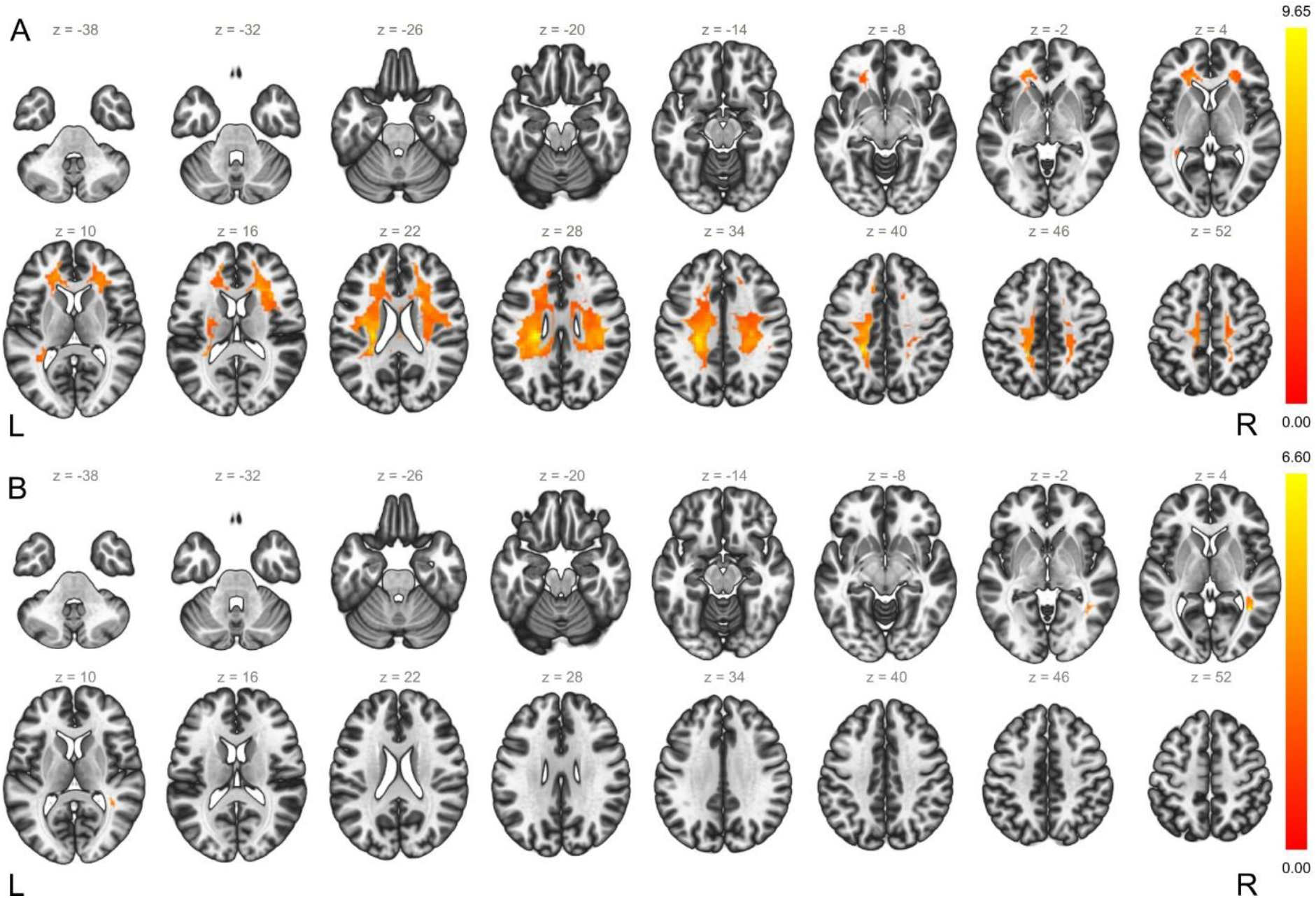
WM Specific ICA. **(A)** Independent component 8. **(B)** Group comparison between VA+ and VA- patients showing a cluster predominantly composed of right superior longitudinal fasciculus.

### 5.2 Regions of Interest Analysis

#### 5.2.1 GM Specific Regions of Interest Analysis

GM specific ROI analysis resulted in a single group of 2 ROI connections (Figure 6A) with significant differences in functional connectivity in the VA+ compared to VA- group (F(2,19) = 16.23, pFDR < 0.01). The 2 connections were between the following resting state networks: 1) posterior division of left superior temporal gyrus (pSTG L) and anterior cerebellar network ROIs, with VA+ group showing increased functional connectivity compared to VA- group; 2) posterior division of right superior temporal gyrus (pSTG R) and posterior cerebellar network ROIs, with VA+ group showing decreased functional connectivity compared to VA- group. As ROI analysis does not provide localization, we performed a post-hoc seed-based analysis using the bilateral pSTG resting networks as seeds to possibly localise regions in the cerebellar networks. We found no significant differences between VA+ and VA- groups after correction for multiple comparisons, however results with uncorrected cluster p-values are reported here to provide some localisation in cerebellar regions. Post-hoc seed based analysis revealed two clusters (Figure 6B): 1) cluster 1 with MNI coordinates of cluster centre (−09, −75, −21) with regions including left cerebellum 6 and left cerebellum crus 1 (F(2,19) = 29.04, p < 0.01 uncorrected); 2) cluster 2 with MNI coordinates of cluster centre (+30, −63, −21) and was composed of right cerebellum 6 (F(2,19) = 16.31, p < 0.05 uncorrected).

**Figure 6.**
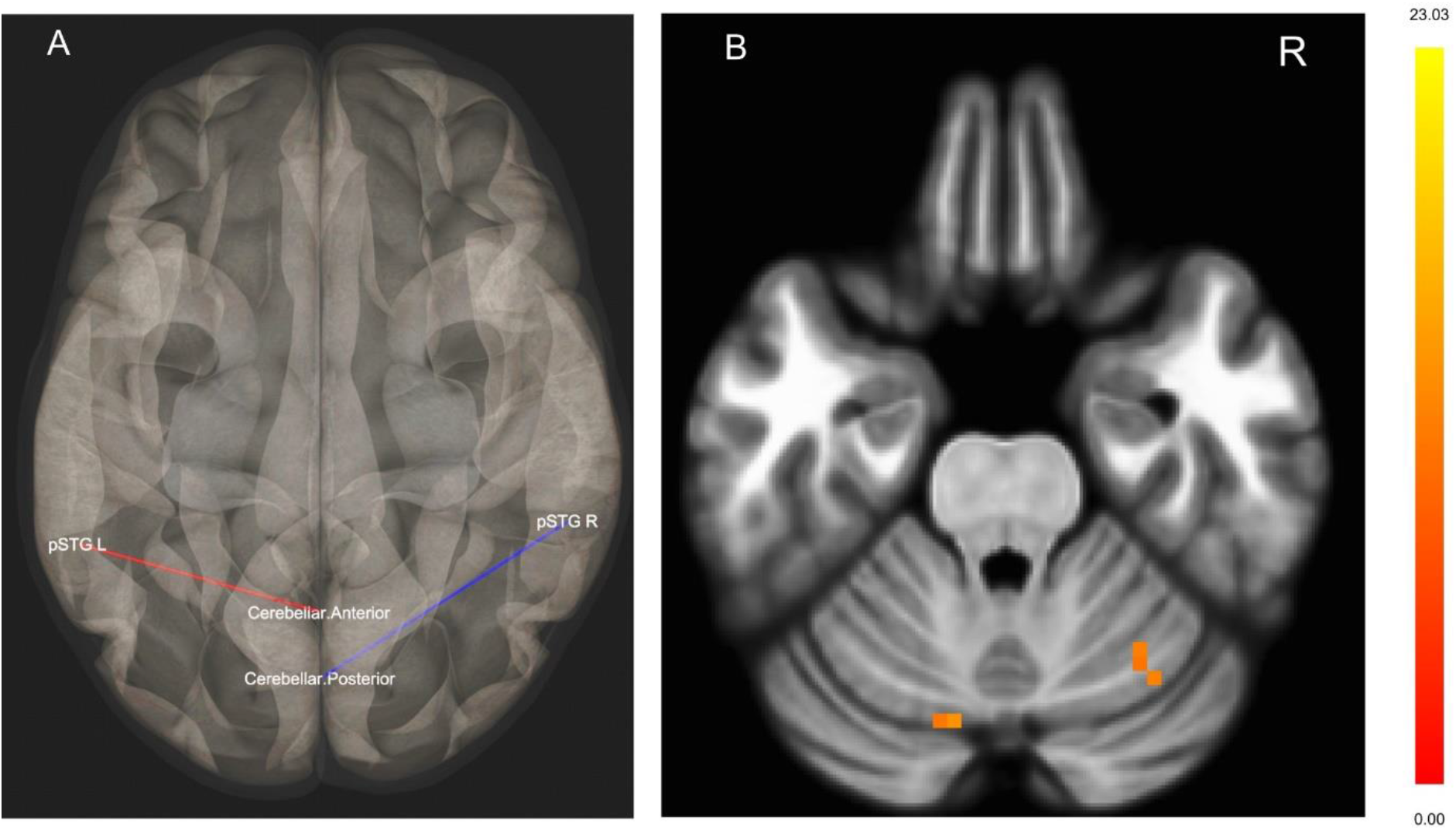
GM Specific ROI Analysis. **(A)** Bilateral connections between superior temporal gyrus and cerebellar resting state ROIs. **(B)** Regions within cerebellum resting state ROIs localised to left cerebellum 6, left cerebellum crus 1, and right cerebellum 6 using bilateral superior temporal gyri as seeds.

#### 5.2.2 WM Specific Regions of Interest Analysis

WM specific ROI analysis resulted in a single group of 6 connections (Figure 7), all with increased functional connectivity in VA+ compared to VA- group (ROI mass = 66.17, pFDR < 0.05). Genu of corpus callosum (GCC), left superior longitudinal fasciculus (SLF), right SLF, left superior corona radiata (SCR), left posterior thalamic radiation (PTR), and left superior fronto-occipital fasciculus (SFOF) were all functionally connected to left anterior corona radiata (ACR), shown in Figure 7.

**Figure 7.**
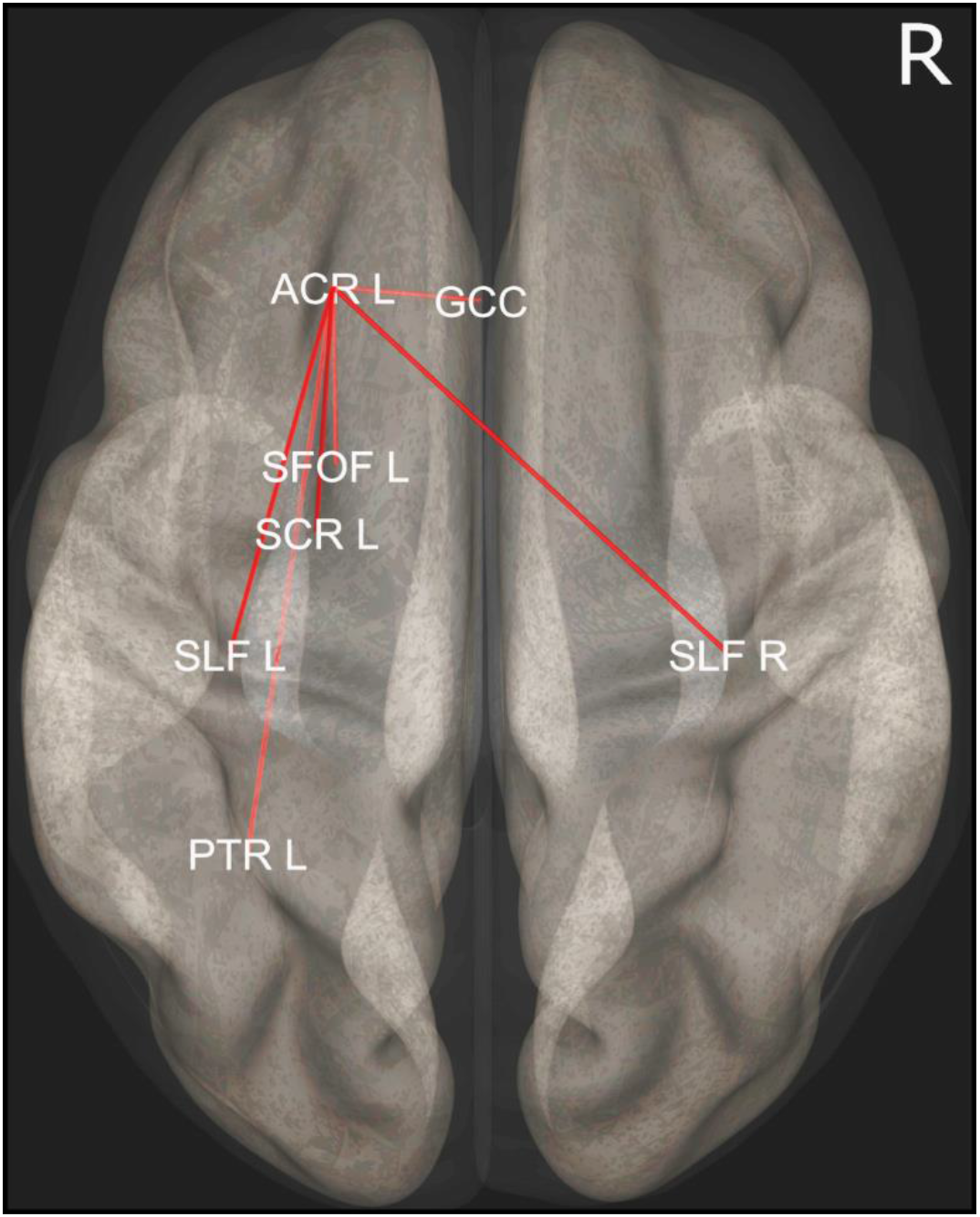
WM Specific ROI Analysis. A group of ROI connections including genu of corpus callosum (GCC), left superior longitudinal fasciculus (SLF), right SLF, left superior corona radiata (SCR), left posterior thalamic radiation (PTR), and left superior fronto-occipital fasciculus (SFOF), all functionally linked to left Anterior Corona Radiata (ACR).

### 5.3 Seed-Based Analysis

#### 5.3.1 Regions with Altered Functional Connections to Vestibular Networks

IC-13, IC-20, and IC-23, determined via GM specific ICA and composed of putative vestibular regions, were selected as seeds for seed-based analysis (supplementary Figure S1). We found that left thalamus (MNI: −12, −27, 12) had increased functional connectivity with the seed networks (F(3,18) = 24.72, pFDR < 0.05) in VA+ group compared to VA- group (Figure 8A). The region was localised to be a part of antero-dorsal thalamus (medial and lateral) using Melbourne Subcortex Atlas (Tian et al., 2020).

**Figure 8.**
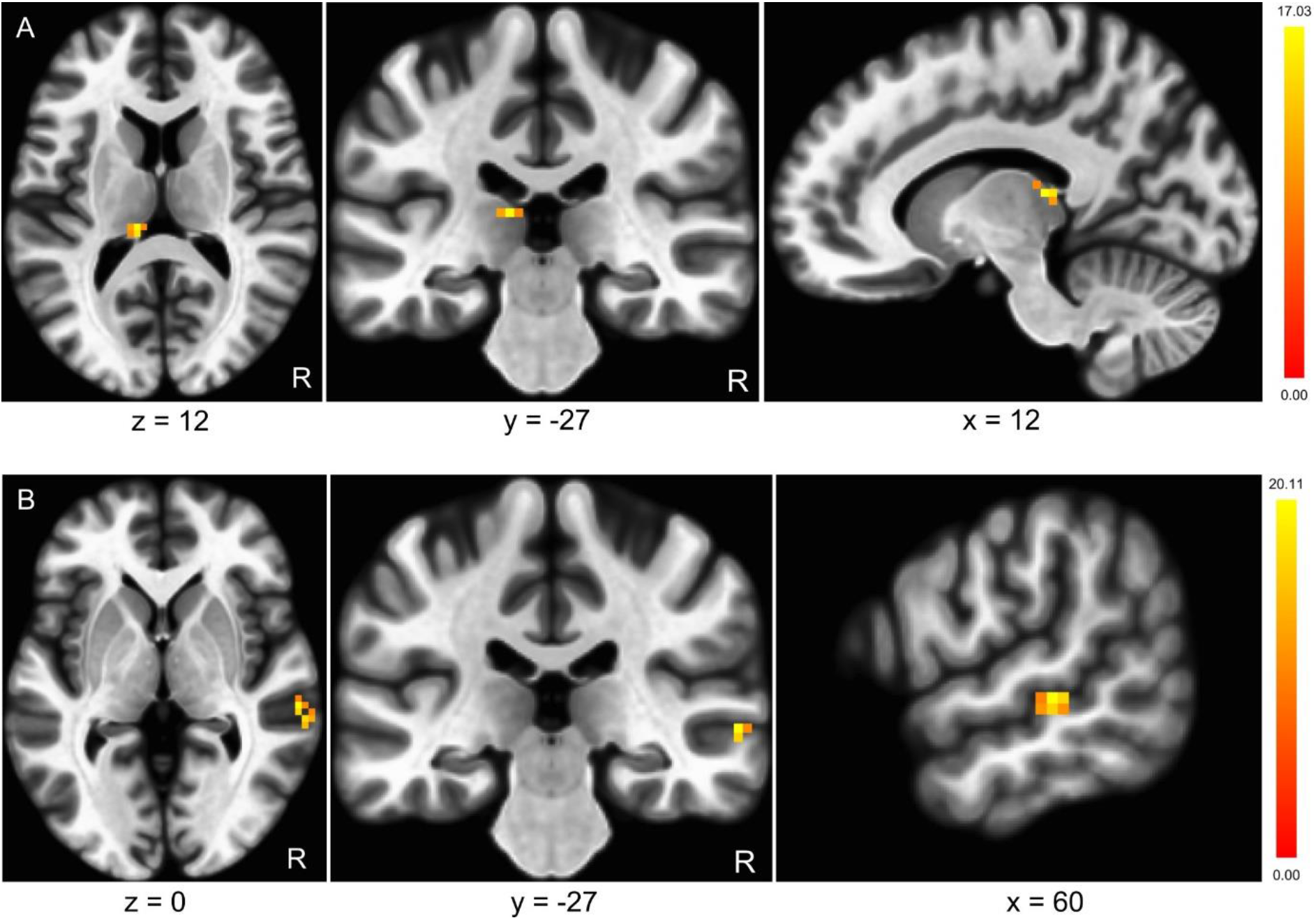
Disconnection between temporal and higher order visual cortices. **(A)** Left thalamus had increased functional connectivity with putative vestibular regions. **(B)** Increased functional connectivity of right mid- and superior-temporal regions when using bilateral lingual gyri as seed.

#### 5.3.2 Disconnection Between Temporal and Higher Order Visual Cortices

Using bilateral lingual gyrus (LG) as seed regions, we found a cluster composed of superior and mid temporal regions with increased functional connectivity in the VA+ compared to the VA- group (F(2,19) = 23.47, pFDR < 0.05). MNI coordinates of cluster centre were (60, −27, 00) and the cluster was composed of superior and mid temporal regions (Figure 8B).

## 6 Discussion

Using DTI imaging, we previously identified that VA was linked to imbalance via damage to right ILF (Calzolari et al., 2021), however, we were unable to identify brain regions specifically explaining VA. Given our a priori hypothesis that vestibular-motion perception is mediated by multiple brain networks, we used resting-state fMRI to identify the functional brain networks linked to vestibular agnosia. We found that vestibular agnosia is linked to: (i) increased functional connectivity in a bilateral white-matter network; (ii) increased functional connectivity in SLF; (iii) a disrupted functional link between regions mapped by ILF (Lingual gyrus (V3v/V4) and mid/superior temporal regions); (iv) a disrupted functional link between the left thalamus and resting state networks containing putative vestibular regions (PIVC, insular, and superior/mid temporal regions).

### 6.1 Vestibular Agnosia and White-Matter fMRI

Using white-matter ICA, we found that vestibular agnosia was linked to a cluster predominantly composed of right superior longitudinal fasciculus (SLF). The cluster also included voxels from right posterior thalamic radiation (PTR), right retrolenticular part of internal capsule (RLIC), and right saggital stratum (SS) (Figure 5B).

Previous electrical stimulation of both right SLF (Spena et al., 2006) and left SLF (Kahane et al., 2003) has shown to induce sensation of self-motion indicating a bilateral coding of self-motion sensations via SLF. We also previously found that a bilateral network mainly composed of the SLF was associated with duration of self-motion sensation following whole body yaw roational accelerations (‘stopping responses’) in the dark in healthy subjects (Nigmatullina et al., 2015). The right retrolenticular internal capsule (RLIC) and right posterior thalamic radiation (PTR) have been linked with impaired balance in mild TBI and elderly adults (Gattu et al., 2016; Rosario et al., 2016) and impaired DTI parameters in RLIC are also associated with impaired spatial navigation (Wu et al., 2016). Since we controlled for the variance explained by balance in our resting-state analysis, our results suggest that the RLIC may function as a vestibular hub, mediating multiple vestibular functions, such as balance, spatial navigation, and vestibular perception of self-motion. Moreover, we have previously shown that right posterior thalamic radiation (PTR) and the right saggital stratum (SS) are linked to vestibular perception of self-motion in patients with impaired balance (Calzolari et al., 2021). Notably the right ILF tract passes through the right SS, and right ILF intra-cortical electrical stimulation has been shown to induce rotational self-motion sensation (Kahane et al., 2003).

While looking at inter-network differences using ROI analysis, the white-matter network found is left lateralized with two interhemispheric connections. One between the left anterior corona radiata (ACR) and right SLF and second a callosal connection between left ACR and genu of corpus callosum (GCC) (Figure 7). Abnormal DTI parameters in the same network of white-matter tracts shown in Figure 7 has previously been reported to be linked to impaired balance in elderly patients with small vessel disease (Rosario et al., 2016), although this study did not assess patients’ ability to perceive self-motion. Notably however, there are multiple accounts of VA in patients with small vessel disease and impaired balance (Chiarovano et al., 2016; Imbaud Genieys, 2007; Seemungal, 2006), thus suggesting the involvement of overlapping albeit distinct networks, mediating multiple vestibular functions.

### 6.2 Vestibular Agnosia and Grey-Matter fMRI

Using ICA we found that a decreased activity, of the right Intra Calcarine Cortex (ICC) in a bilateral mid-temporal resting-state network, and an overactivation of left frontal pole in rostral prefrontal cortical network, were linked to vestibular agnosia.

Mid-temporal gyrus is linked with rotational sensation of self-motion induced by intra-cortical electrical stimulation (Kahane et al., 2003) and during non-invasive galvanic stimulation (Della-Justina et al., 2014). Additionally, the calcarine region is often shown to be linked with visual motion and visuo-vestibular interaction (Cha & Chakrapani, 2015; Della-Justina et al., 2014). Since the ILF connects the calcarine and temporal regions (Latini et al., 2017; Panesar et al., 2018), the deactivation of the ICC within a bilateral mid-temporal resting-state network might be due to the damage to the ILF, which we previously showed in this cohort via DTI (Calzolari et al., 2021).

Left frontal pole region is located within Rostral prefrontal cortex (RPFC) resting state network, which is well established as a region associated with the integration of sensory information (Stocco et al., 2012). In contrast to the integration hypotheses, it’s possible that RPFC may also work by perceptual ‘ignition’ mechanism, in which perception is triggered after enough evidence is accumulated (Del Cul et al., 2007, 2009).

While looking at the inter-network differences using ROI approach, we found bilateral functional connections between the superior temporal gyri and cerebellar resting-state networks. There was a relative increase in activation of left hemispheric connections and a decrease in right hemisphere connections, for VA+ patients when compared to VA- patients. Since resting-state networks do not provide any localization of regions per se, we localized cerebellar regions using a seed-based analysis. There were no statistically significant differences, however, uncorrected findings (for multiple comparisons and hence require a conservative interpretation) showed bilateral functional connections from temporal gyri localized to the left cerebellum VI/Crus I, previously shown to activate in response to vestibular stimulation (Della-Justina et al., 2014), and right cerebellum VI. Taken together, our results point to the existence of bilateral functional connections from temporal gyri to the cerebellum.

### 6.3 Altered Connectivity of Vestibular Networks Linked to Vestibular Agnosia

We adopted a network-based approach and used the resting state networks composed of core vestibular regions (PIVC and PIC (Wirth et al., 2018)) in a seed-based analysis. We found that patients with vestibular agnosia (VA+) had an overactivated functional link between the left thalamus and the vestibular resting-state networks.

Notably, the MNI coordinates of the left thalamus localized to the medial and lateral parts of the antero-dorsal thalamus, of potential relevance since the thalamus is an important vestibular structure and is considered a ‘gateway’ for vestibular information (Dieterich et al., 2005) with bidirectional projections linking brainstem and cortical vestibular regions (Kirsch et al., 2016). Previous tractography studies have shown that PIVC and PIC have structural connections with the thalamus in each hemisphere (Kirsch et al., 2016; Wirth et al., 2018). Most of the previous studies have reported posterolateral parts of thalamus to be important centers for vestibular processing including ventral posterior lateral nucleus (VPL), ventral posterior medial (VPM) nucleus, lateral posterior (LP) nucleus, and ventral lateral posterior or ventro-intermediate nucleus (VLP/Vim) (Baier et al., 2016, 2017; Dieterich et al., 2005; Dieterich & Brandt, 1993). However, it is important to note that most of these findings have been reported for visual tilt perception tasks and do not provide much information about self-motion perception processing.

### 6.4 Disconnection Between Higher Order Visual Cortices and Temporal Regions

We found that VA+ subjects had a disruption of functional link between right lingual gyrus and right mid-temporal gyrus indicating disrupted functional projections from higher order visual cortex in these patients. Given our previous report of ILF white matter microstructural disruption in VA patients (Calzolari et al., 2021), and the known connectivity of the ILF linking the temporal and lingual gyri (Latini et al., 2017; Panesar et al., 2018), this pattern of disruption was perhaps predictable.

The lingual gyrus appears to be an important vestibular processing hub that is also involved in visuo-vestibular motion processing (Della-Justina et al., 2014; Roberts et al., 2017). The lingual gyrus has been shown to have a specific role in vestibular processing (Della-Justina et al., 2014; Roberts et al., 2017). Indeed intra-cortical electrical stimulation of the lingual gyrus evoked a yaw-plane sensation of self-motion (Kahane et al., 2003). Hence, in addition to previous data implicating the lingual gyrus in motion processing, our data further suggest that a right hemisphere network involving the lingual gyrus and mid-temporal regions connected via the right ILF, is involved in self-motion processing.

### 6.5 Limitations

Vestibular agnosia (VA), while clinically apparent, can currently only be objectively identified via self-motion perception testing in a rotatory-chair in dark. Moreover, VA is classified based on a continuous parameter i.e. sensory thresholds of motion perception, and thus mild or borderline cases may or may not be classified as having VA. Another limitation of the study is the small sample size. This was partly due to a limited number of vestibular agnosia patients, which is unavoidable considering that prevalence of VA is not yet known and patients are classified post-hoc based on objective laboratory testing. However, similar and even smaller sample sizes have previously been used in TBI cohorts to adequately identify functional brain differences (De Simoni et al., 2016; Sours et al., 2015; Woytowicz et al., 2018). VA is known to impact elderly people and neurodegeneration patients, however an advantage of our patient group was their relative young age and good premorbid health excluding incipient neurodegenerative disease (given our stringent exclusion criteria), hence the findings reported herein were overwhelmingly related to the acute TBI and not to other chronic underlying disease.

### 6.6 Conclusion

Our data provide the first evidence linking resting-state functional networks to vestibular agnosia – impaired self-motion perception. More specifically, we show that self-motion perception is mediated via bihemispheric brain networks, composed of posterior vestibular regions involved in sensory processing and anterior regions involved in sensory integration and perceptual ignition.

## Declaration of Competing Interests

The authors declare that they have no competing interests.

## Acknowledgements

We would like to thank our patient and healthy volunteers for their participation and for helping us improve the traumatic brain injury patients’ care. We are also grateful to the major trauma ward teams at St Mary’s Hospital and King’s College Hospital London for their help with recruitment and assessment. We are also very grateful to The Imperial Health Charity who provided important kickstarter funding that enabled us to obtain research council funding for this project.

## Funding

The Medical Research Council (MRC). The Imperial Health Charity, The NIHR Imperial Biomedical Research Centre, NIHR Clinical Doctoral Research Fellowship, The US Department of Defense - Congressionally Directed Medical Research Program (CDMRP). The Jon Moulton Charity Trust.

## Data and code availability statements

Raw data that support the findings of this study are available from the corresponding author, upon reasonable request. The request would require a formal data sharing agreement, approval from the requesting researcher’s local ethics committee, a formal project outline, and discussion about authorship on any research output from the shared data.

## Author Contributions

**Zaeem Hadi**: Investigation, Methodology, Software, Formal analysis, Visualization, Writing - Original Draft. **Yuscah Pondeca**: Investigation, Methodology, Writing – review and editing. **Elena Calzolari**: Investigation, Methodology, Writing – review and editing. **Mariya Chepisheva**: Investigation, Writing – review and editing. **Rebecca M Smith**: Investigation, Methodology, Writing – review and editing. **Mohammad Mahmud**: Investigation, Methodology, Writing – review and editing. **Heiko Rust**: Investigation, Methodology, Writing – review and editing. **David J Sharp**: Supervision, Writing – review and editing. **Barry M Seemungal**: Conceptualization, Investigation, Project administration, Funding acquisition, Resources, Supervision, Writing – review and editing.

## Notes

### Competing Interest Statement

The authors have declared no competing interest.

## References

Baier, B., Conrad, J., Stephan, T., Kirsch, V., Vogt, T., Wilting, J., Müller-Forell, W., & Dieterich, M. (2016). Vestibular thalamus Two distinct graviceptive pathways. Neurology, 86(2), 134–140. https://doi.org/10.1212/WNL.0000000000002238

Baier, B., Vogt, T., Rohde, F., Cuvenhaus, H., Conrad, J., & Dieterich, M. (2017). Deep brain stimulation of the nucleus ventralis intermedius: a thalamic site of graviceptive modulation. Brain Structure and Function, 222(1), 645–650. https://doi.org/10.1007/s00429-015-1157-x

Behzadi, Y., Restom, K., Liau, J., & Liu, T. T. (2007). A component based noise correction method (CompCor) for BOLD and perfusion based fMRI. NeuroImage, 37(1), 90–101. https://doi.org/10.1016/j.neuroimage.2007.04.042

Branco, P., Seixas, D., & Castro, S. L. (2020). Mapping language with resting-state functional magnetic resonance imaging: A study on the functional profile of the language network. Human Brain Mapping, 41(2), 545–560. https://doi.org/10.1002/HBM.24821

Branco, P., Seixas, D., Deprez, S., Kovacs, S., Peeters, R., Castro, S. L., & Sunaert, S. (2016). Resting-State Functional Magnetic Resonance Imaging for Language Preoperative Planning. Frontiers in Human Neuroscience, 0(FEB2016), 11. https://doi.org/10.3389/FNHUM.2016.00011

Bullmore, E. T., Suckling, J., Overmeyer, S., Rabe-Hesketh, S., Taylor, E., & Brammer, M. J. (1999). Global, voxel, and cluster tests, by theory and permutation, for a difference between two groups of structural mr images of the brain. IEEE Transactions on Medical Imaging, 18(1), 32–42. https://doi.org/10.1109/42.750253

Calzolari, E., Chepisheva, M., Smith, R. M., Mahmud, M., Hellyer, P. J., Tahtis, V., Arshad, Q., Jolly, A., Wilson, M., Rust, H., Sharp, D. J., & Seemungal, B. M. (2021). Vestibular agnosia in traumatic brain injury and its link to imbalance. Brain: A Journal of Neurology, 144(1), 128–143. https://doi.org/10.1093/brain/awaa386

Cha, Y.-H., & Chakrapani, S. (2015). Voxel Based Morphometry Alterations in Mal de Debarquement Syndrome. PLOS ONE, 10(8), e0135021. https://doi.org/10.1371/journal.pone.0135021

Chiarovano, E., Vidal, P. P., Magnani, C., Lamas, G., Curthoys, I. S., & de Waele, C. (2016). Absence of rotation perception during warm water caloric irrigation in some seniors with postural instability. Frontiers in Neurology, 7(JAN). https://doi.org/10.3389/fneur.2016.00004

De Simoni, S., Grover, P. J., Jenkins, P. O., Honeyfield, L., Quest, R. A., Ross, E., Scott, G., Wilson, M. H., Majewska, P., Waldman, A. D., Patel, M. C., & Sharp, D. J. (2016). Disconnection between the default mode network and medial temporal lobes in post-traumatic amnesia. Brain, 139(12), 3137. https://doi.org/10.1093/BRAIN/AWW241

Del Cul, A., Baillet, S., & Dehaene, S. (2007). Brain dynamics underlying the nonlinear threshold for access to consciousness. PLoS Biology, 5(10), 2408–2423. https://doi.org/10.1371/journal.pbio.0050260

Del Cul, A., Dehaene, S., Reyes, P., Bravo, E., & Slachevsky, A. (2009). Causal role of prefrontal cortex in the threshold for access to consciousness. Brain, 132(9), 2531–2540. https://doi.org/10.1093/brain/awp111

Della-Justina, H. M., Gamba, H. R., Lukasova, K., Nucci-da-Silva, M. P., Winkler, A. M., & Amaro, E. (2014). Interaction of brain areas of visual and vestibular simultaneous activity with fMRI. Experimental Brain Research, 233(1), 237–252. https://doi.org/10.1007/s00221-014-4107-6

Dieterich, M., Bartenstein, P., Spiegel, S., Bense, S., Schwaiger, M., & Brandt, T. (2005). Thalamic infarctions cause side-specific suppression of vestibular cortex activations. Brain, 128(9), 2052–2067. https://doi.org/10.1093/brain/awh551

Dieterich, M., & Brandt, T. (1993). Thalamic infarctions: Differential effects on vestibular function in the roll plane (35 patients). Neurology, 43(9), 1732–1740. https://doi.org/10.1212/wnl.43.9.1732

Ding, Z., Newton, A. T., Xu, R., Anderson, A. W., Morgan, V. L., & Gore, J. C. (2013). Spatio-temporal correlation tensors reveal functional structure in human brain. PLoS ONE, 8(12), 82107. https://doi.org/10.1371/journal.pone.0082107

Ding, Z., Xu, R., Bailey, S. K., Wu, T. L., Morgan, V. L., Cutting, L. E., Anderson, A. W., & Gore, J. C. (2016). Visualizing functional pathways in the human brain using correlation tensors and magnetic resonance imaging. Magnetic Resonance Imaging, 34(1), 8–17. https://doi.org/10.1016/j.mri.2015.10.003

Dotson, N. M., Salazar, R. F., & Gray, C. M. (2014). Frontoparietal Correlation Dynamics Reveal Interplay between Integration and Segregation during Visual Working Memory. Journal of Neuroscience, 34(41), 13600–13613. https://doi.org/10.1523/JNEUROSCI.1961-14.2014

Dovern, A., Fink, G. R., Fromme, A. C. B., Wohlschläger, A. M., Weiss, P. H., & Riedl, V. (2012). Intrinsic Network Connectivity Reflects Consistency of Synesthetic Experiences. Journal of Neuroscience, 32(22), 7614–7621. https://doi.org/10.1523/JNEUROSCI.5401-11.2012

Eklund, A., Knutsson, H., & Nichols, T. E. (2019). Cluster failure revisited: Impact of first level design and physiological noise on cluster false positive rates. Human Brain Mapping, 40(7), 2017–2032. https://doi.org/10.1002/hbm.24350

Ester, E. F., Sutterer, D. W., Serences, J. T., & Awh, E. (2016). Feature-Selective Attentional Modulations in Human Frontoparietal Cortex. Journal of Neuroscience, 36(31), 8188–8199. https://doi.org/10.1523/JNEUROSCI.3935-15.2016

Fiebelkorn, I. C., Pinsk, M. A., & Kastner, S. (2018). A Dynamic Interplay within the Frontoparietal Network Underlies Rhythmic Spatial Attention. Neuron, 99(4), 842–853.e8. https://doi.org/10.1016/J.NEURON.2018.07.038

Fraser, L. M., Stevens, M. T., Beyea, S. D., & D’Arcy, R. C. N. (2012). White versus gray matter: fMRI hemodynamic responses show similar characteristics, but differ in peak amplitude. BMC Neuroscience, 13(1), 91. https://doi.org/10.1186/1471-2202-13-91

Galati, G., Committeri, G., Sanes, J. N., & Pizzamiglio, L. (2001). Spatial coding of visual and somatic sensory information in body-centred coordinates. European Journal of Neuroscience, 14(4), 737–746. https://doi.org/10.1046/J.0953-816X.2001.01674.X

Gattu, R., Akin, F. W., Cacace, A. T., Hall, C. D., Murnane, O. D., & Haacke, E. M. (2016). Vestibular, balance, microvascular and white matter neuroimaging characteristics of blast injuries and mild traumatic brain injury: Four case reports. Brain Injury, 30(12), 1501–1514. https://doi.org/10.1080/02699052.2016.1219056

Huang, Y., Bailey, S. K., Wang, P., Cutting, L. E., Gore, J. C., & Ding, Z. (2018). Voxel-wise detection of functional networks in white matter. NeuroImage, 183, 544–552. https://doi.org/10.1016/j.neuroimage.2018.08.049

Huang, Y., Yang, Y., Hao, L., Hu, X., Wang, P., Ding, Z., Gao, J. H., & Gore, J. C. (2020). Detection of functional networks within white matter using independent component analysis. NeuroImage, 222, 117278. https://doi.org/10.1016/j.neuroimage.2020.117278

Imbaud Genieys, S. (2007). Vertigo, dizziness and falls in the elderly. Annales d’Oto-Laryngologie et de Chirurgie Cervico-Faciale, 124(4), 189–196. https://doi.org/10.1016/j.aorl.2007.04.003

Jafri, M. J., Pearlson, G. D., Stevens, M., & Calhoun, V. D. (2008). A method for functional network connectivity among spatially independent resting-state components in schizophrenia. NeuroImage, 39(4), 1666–1681. https://doi.org/10.1016/j.neuroimage.2007.11.001

Kahane, P., Hoffmann, D., Minotti, L., & Berthoz, A. (2003). Reappraisal of the Human Vestibular Cortex by Cortical Electrical Stimulation Study. Annals of Neurology, 54(5), 615–624. https://doi.org/10.1002/ana.10726

Kaski, D., Quadir, S., Nigmatullina, Y., Malhotra, P. A., Bronstein, A. M., & Seemungal, B. M. (2016). Temporoparietal encoding of space and time during vestibular-guided orientation. Brain, 139(2), 392–403. https://doi.org/10.1093/brain/awv370

Kirsch, V., Keeser, D., Hergenroeder, T., Erat, O., Ertl-Wagner, B., Brandt, T., & Dieterich, M. (2016). Structural and functional connectivity mapping of the vestibular circuitry from human brainstem to cortex. Brain Structure and Function, 221(3), 1291–1308. https://doi.org/10.1007/s00429-014-0971-x

Latini, F., Mårtensson, J., Larsson, E. M., Fredrikson, M., Åhs, F., Hjortberg, M., Aldskogius, H., & Ryttlefors, M. (2017). Segmentation of the inferior longitudinal fasciculus in the human brain: A white matter dissection and diffusion tensor tractography study. Brain Research, 1675, 102–115. https://doi.org/10.1016/j.brainres.2017.09.005

Li, M., Gao, Y., Gao, F., Anderson, A. W., Ding, Z., & Gore, J. C. (2020). Functional engagement of white matter in resting-state brain networks. NeuroImage, 220, 117096. https://doi.org/10.1016/j.neuroimage.2020.117096

Marussich, L., Lu, K. H., Wen, H., & Liu, Z. (2017). Mapping white-matter functional organization at rest and during naturalistic visual perception. NeuroImage, 146, 1128–1141. https://doi.org/10.1016/j.neuroimage.2016.10.005

Mori, S., Oishi, K., Jiang, H., Jiang, L., Li, X., Akhter, K., Hua, K., Faria, A. V., Mahmood, A., Woods, R., Toga, A. W., Pike, G. B., Neto, P. R., Evans, A., Zhang, J., Huang, H., Miller, M. I., van Zijl, P., & Mazziotta, J. (2008). Stereotaxic white matter atlas based on diffusion tensor imaging in an ICBM template. NeuroImage, 40(2), 570–582. https://doi.org/10.1016/j.neuroimage.2007.12.035

Murphy, K., Birn, R. M., Handwerker, D. A., Jones, T. B., & Bandettini, P. A. (2009). The impact of global signal regression on resting state correlations: Are anti-correlated networks introduced? NeuroImage, 44(3), 893–905. https://doi.org/10.1016/j.neuroimage.2008.09.036

Nichols, T. E. (2012). Multiple testing corrections, nonparametric methods, and random field theory. NeuroImage, 62(2), 811–815. https://doi.org/10.1016/J.NEUROIMAGE.2012.04.014

Nigmatullina, Y., Hellyer, P. J., Nachev, P., Sharp, D. J., & Seemungal, B. M. (2015). The neuroanatomical correlates of training-related perceptuo-reflex uncoupling in dancers. Cerebral Cortex, 25(2), 554–562. https://doi.org/10.1093/cercor/bht266

Ogawa, S., Tank, D. W., Menon, R., Ellermann, J. M., Kim, S. G., Merkle, H., & Ugurbil, K. (1992). Intrinsic signal changes accompanying sensory stimulation: Functional brain mapping with magnetic resonance imaging. Proceedings of the National Academy of Sciences of the United States of America, 89(13), 5951–5955. https://doi.org/10.1073/pnas.89.13.5951

Oishi, K., Zilles, K., Amunts, K., Faria, A., Jiang, H., Li, X., Akhter, K., Hua, K., Woods, R., Toga, A. W., Pike, G. B., Rosa-Neto, P., Evans, A., Zhang, J., Huang, H., Miller, M. I., van Zijl, P. C. M., Mazziotta, J., & Mori, S. (2008). Human brain white matter atlas: Identification and assignment of common anatomical structures in superficial white matter. NeuroImage, 43(3), 447–457. https://doi.org/10.1016/j.neuroimage.2008.07.009

Panesar, S. S., Yeh, F. C., Jacquesson, T., Hula, W., & Fernandez-Miranda, J. C. (2018). A quantitative tractography study into the connectivity segmentation laterality of the human inferior longitudinal fasciculus. Frontiers in Neuroanatomy, 12. https://doi.org/10.3389/fnana.2018.00047

Peer, M., Nitzan, M., Bick, A. S., Levin, N., & Arzy, S. (2017). Evidence for functional networks within the human brain’s white matter. Journal of Neuroscience, 37(27), 6394–6407. https://doi.org/10.1523/JNEUROSCI.3872-16.2017

Power, J. D., Barnes, K. A., Snyder, A. Z., Schlaggar, B. L., & Petersen, S. E. (2012). Spurious but systematic correlations in functional connectivity MRI networks arise from subject motion. NeuroImage, 59(3), 2142–2154. https://doi.org/10.1016/j.neuroimage.2011.10.018

Roberts, R. E., Ahmad, H., Arshad, Q., Patel, M., Dima, D., Leech, R., Seemungal, B. M., Sharp, D. J., & Bronstein, A. M. (2017). Functional neuroimaging of visuo-vestibular interaction. Brain Structure and Function, 222(5), 2329–2343. https://doi.org/10.1007/s00429-016-1344-4

Rosario, B. L., Rosso, A. L., Aizenstein, H. J., Harris, T., Newman, A. B., Satterfield, S., Studenski, S. A., Yaffe, K., & Rosano, C. (2016). Cerebral white matter and slow gait: Contribution of hyperintensities and normal-appearing parenchyma. Journals of Gerontology - Series A Biological Sciences and Medical Sciences, 71(7), 968–973. https://doi.org/10.1093/gerona/glv224

Seemungal, B. M. (2006). The mechanisms and loci of human vestibular perception [University of London]. https://discovery.ucl.ac.uk/id/eprint/1445054/

Seemungal, B. M., Gunaratne, I. A., Fleming, I. O., Gresty, M. A., & Bronstein, A. M. (2004). Perceptual and nystagmic thresholds of vestibular function in yaw. Journal of Vestibular Research, 14(6), 461–466.

Seemungal, B. M., Rizzo, V., Gresty, M. A., Rothwell, J. C., & Bronstein, A. M. (2008). Posterior parietal rTMS disrupts human Path Integration during a vestibular navigation task. Neuroscience Letters, 437(2), 88–92. https://doi.org/10.1016/j.neulet.2008.03.067

Sours, C., Zhuo, J., Roys, S., Shanmuganathan, K., & Gullapalli, R. P. (2015). Disruptions in Resting State Functional Connectivity and Cerebral Blood Flow in Mild Traumatic Brain Injury Patients. PloS One, 10(8). https://doi.org/10.1371/JOURNAL.PONE.0134019

Spena, G., Gatignol, P., Capelle, L., & Duffau, H. (2006). Superior longitudinal fasciculus subserves vestibular network in humans. NeuroReport, 17(13), 1403–1406. https://doi.org/10.1097/01.wnr.0000223385.49919.61

Stocco, A., Lebiere, C., O’Reilly, R. C., & Anderson, J. R. (2012). Distinct contributions of the caudate nucleus, rostral prefrontal cortex, and parietal cortex to the execution of instructed tasks. Cognitive, Affective and Behavioral Neuroscience, 12(4), 611–628. https://doi.org/10.3758/s13415-012-0117-7

Tian, Y., Margulies, D. S., Breakspear, M., & Zalesky, A. (2020). Topographic organization of the human subcortex unveiled with functional connectivity gradients. Nature Neuroscience, 23(11), 1421–1432. https://doi.org/10.1038/s41593-020-00711-6

Tie, Y., Rigolo, L., Norton, I. H., Huang, R. Y., Wu, W., Orringer, D., Mukundan, S., & Golby, A. J. (2014). Defining language networks from resting-state fMRI for surgical planning—a feasibility study. Human Brain Mapping, 35(3), 1018–1030. https://doi.org/10.1002/HBM.22231

Whitfield-Gabrieli, S., & Nieto-Castanon, A. (2012). Conn: A Functional Connectivity Toolbox for Correlated and Anticorrelated Brain Networks. Brain Connectivity, 2(3), 125–141. https://doi.org/10.1089/brain.2012.0073

Wirth, A. M., Frank, S. M., Greenlee, M. W., & Beer, A. L. (2018). White Matter Connectivity of the Visual-Vestibular Cortex Examined by Diffusion-Weighted Imaging. Brain Connectivity, 8(4), 235–244. https://doi.org/10.1089/brain.2017.0544

Woytowicz, E. J., Sours, C., Gullapalli, R. P., Rosenberg, J., & Westlake, K. P. (2018). Modulation of working memory load distinguishes individuals with and without balance impairments following mild traumatic brain injury. Brain Injury, 32(2), 191–199. https://doi.org/10.1080/02699052.2017.1403045

Wu, Y. F., Wu, W. B., Liu, Q. P., He, W. W., Ding, H., Nedelska, Z., Hort, J., Zhang, B., & Xu, Y. (2016). Presence of lacunar infarctions is associated with the spatial navigation impairment in patients with mild cognitive impairment: A DTI study. Oncotarget, 7(48), 78310–78319. https://doi.org/10.18632/oncotarget.13409

Yousif, N., Bhatt, H., Bain, P. G., Nandi, D., & Seemungal, B. M. (2016). The effect of pedunculopontine nucleus deep brain stimulation on postural sway and vestibular perception. European Journal of Neurology, 23(3), 668–670. https://doi.org/10.1111/ene.12947

Yushkevich, P. A., Piven, J., Hazlett, H. C., Smith, R. G., Ho, S., Gee, J. C., & Gerig, G. (2006). User-guided 3D active contour segmentation of anatomical structures: Significantly improved efficiency and reliability. NeuroImage, 31(3), 1116–1128. https://doi.org/10.1016/j.neuroimage.2006.01.015

Zhang, H., Nichols, T. E., & Johnson, T. D. (2009). Cluster mass inference via random field theory. NeuroImage, 44(1), 51–61. https://doi.org/10.1016/J.NEUROIMAGE.2008.08.017

Zhu, D. C., Covassin, T., Nogle, S., Doyle, S., Russell, D., Pearson, R. L., Monroe, J., Liszewski, C. M., DeMarco, J. K., & Kaufman, D. I. (2015). A potential biomarker in sports-related concussion: brain functional connectivity alteration of the default-mode network measured with longitudinal resting-state fMRI over thirty days. Journal of Neurotrauma, 32(5), 327–341. https://doi.org/10.1089/NEU.2014.3413

