## Supplementary Figure S1 for "The human brain networks mediating the vestibular sensation of self-motion"

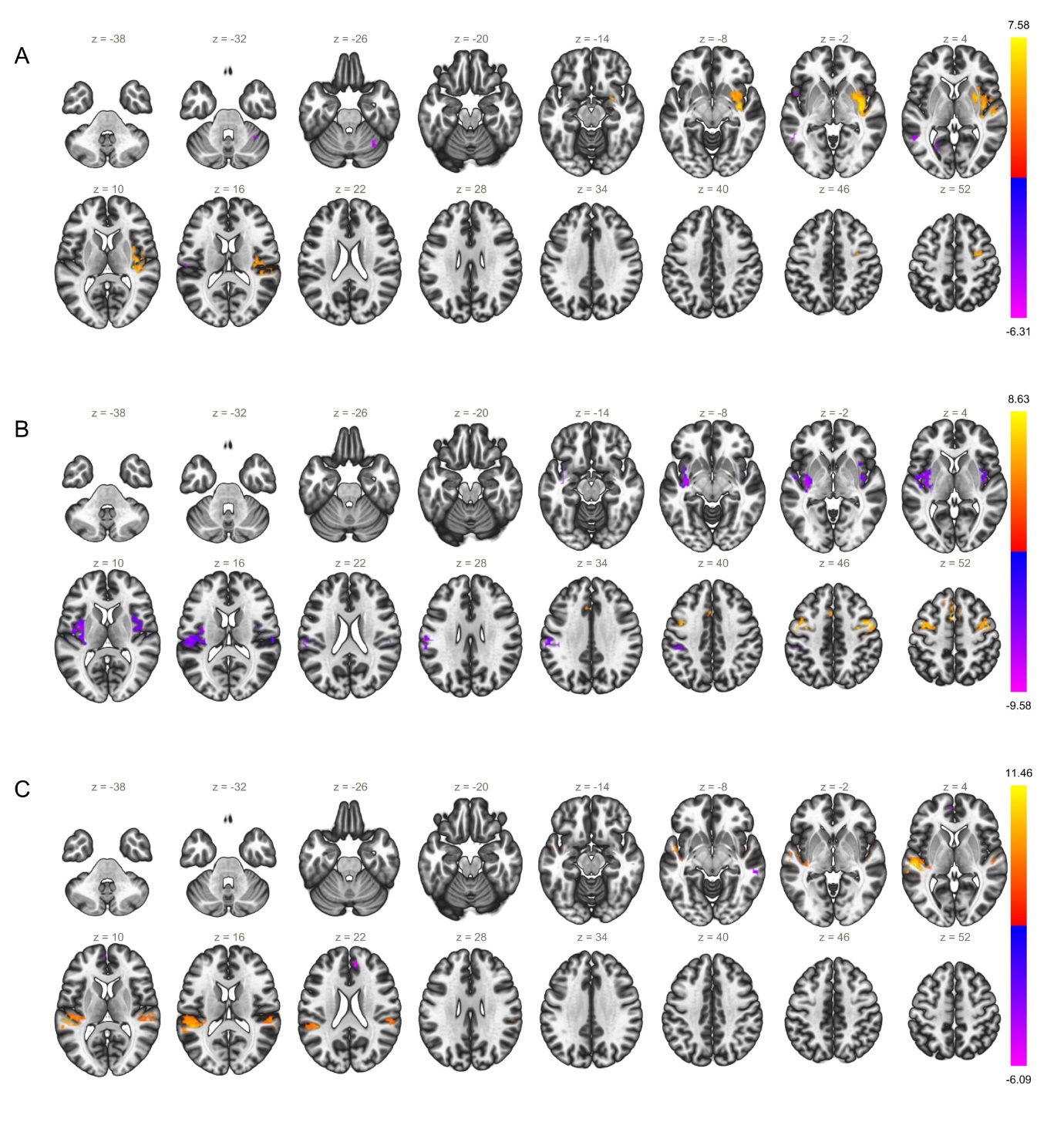

Figure S1 **A.** IC-13: Containing voxels from right insular cortex, right parietal operculum, mid temporal gyrus, and cerebellum 6. **B.** IC-20: Containing voxels from left insular cortex, right and left parietal operculum. **C.** IC-23: Containing voxels from right and left parietal operculum, superior and mid temporal gyrus.
